# A non-catalytic function of carbonic anhydrase IX contributes to the glycolytic phenotype and pH regulation in human breast cancer cells

**DOI:** 10.1101/505644

**Authors:** Mam Y. Mboge, Zhijuan Chen, Daniel Khokhar, Alyssa Wolff, Lingbao Ai, Coy D. Heldermon, Murat Bozdag, Fabrizio Carta, Claudiu T. Supuran, Kevin D. Brown, Robert McKenna, Christopher J. Frost, Susan C. Frost

## Abstract

The most aggressive and invasive tumor cells often reside in hypoxic microenvironments and rely heavily on rapid anaerobic glycolysis for energy production. This switch from oxidative phosphorylation to glycolysis, along with up-regulation of the glucose transport system, significantly increases the release of lactic acid from cells into the tumor microenvironment. Excess lactate and proton excretion exacerbate extracellular acidification to which cancer cells, but not normal cells, adapt. We have hypothesized that carbonic anhydrases (CAs) play a role in stabilizing both intracellular and extracellular pH to favor cancer progression and metastasis. Here we show that proton efflux (acidification) using the glycolytic rate assay is dependent on both extracellular pH (pH_e_) and CA IX expression. Yet, isoform selective sulfonamide-based inhibitors of CA IX did not alter proton flux, which suggests that the catalytic activity of CA IX is not necessary for this regulation. Other investigators have suggested the CA IX cooperates with the MCT transport family to excrete protons. To test this possibility, we examined the expression patterns of selected ion transporters and show that members of this family are differentially expressed within the molecular subtypes of breast cancer. The most aggressive form of breast cancer, triple negative breast cancer (TNBC), appears to coordinately express the monocarboxylate transporter 4 (MCT4) and carbonic anhydrase IX (CA IX). This supports a possible mechanism that utilizes the intramolecular H^+^ shuttle system in CA IX to facilitate proton efflux through MCT4.

## Introduction

The tumor microenvironment provides metabolic challenges to tumor cells (1). Intermittent hypoxia, induced by alterations in blood flow, changes expression of genes involved in glucose metabolism through the transcription factor, hypoxia inducible factor 1 (HIF1) (2–4). Cells switch from dependence on oxidative phosphorylation to accelerated glycolysis for their energy production (4). This glycolytic switch increases the production of lactic acid reducing extracellular pH (pH_e_) (5). Ultimately, this condition becomes imprinted so that it is maintained even in the presence of oxygen, which was described by Otto Warburg nearly 100 years ago and coined the “Warburg Effect” (6–8). Cancer cells that adapt to these conditions proliferate rapidly and are resistant to both radiation and chemotherapy (4). Normal cells in this environment become apoptotic and suffer cell death. This leads to increased space in which cancer cells expand. Thus, the glycolytic phenotype is advantageous for cancer progression and metastasis (3, 4, 8, 9).

To accommodate the increasing concentration of lactic acid inside cells, it has been proposed that proton-specific transporters are induced in cancer cells to normalize intracellular pH (pH_i_) (5). Indeed, it has been shown that pH_i_ in the cytoplasm of tumor cells (pH 7.2-7.4) is similar to that of normal cells (10, 11). However, the pH_e_ surrounding tumor cells is strongly acidic (pH 6.5-6.8) (11). One of the key players in maintaining the relationship between pH_i_ and pH_e_ in tumor cells is thought to be the monocarboxylate transporters (MCT1 and MCT4) (12, 13). These transporters co-transport lactate with a proton. Other proteins that may participate in pH regulation include the Na^+^/H^+^ exchanger (NHE1), the Na^+^/HCO_3_^−^ co-transporter (NBCn1) and the vacuolar ATPase (vATPase) (12, 14–18). Previously published studies performed using glycolytic-deficient cells showed that CO_2_ might also contribute to acidification of the tumor microenvironment (19, 20). CO_2_ is itself a weak acid oxide and can react with water to form HCO_3_^−^ and a H^+^. This process occurs naturally but is enhanced by as much as 1000-fold by carbonic anhydrase family (CAs). CAs are a group of metalloenzymes that consists of five distinct classes, including the α-CAs which is the only class found in humans (21, 22). As shown below, the CA isoforms catalyze a reversible reaction (Overall) and have the potential to produce CO_2_, HCO_3_^−^, and protons to create the pH differential across the plasma membrane of cancer cells. In the first of two independent stages, CO_2_ undergoes hydration within the active site to form a Zn-bound bicarbonate, which is subsequently exchanged for H_2_O releasing the bicarbonate (Equation 1). In the second stage, the Zn-bound water is deprotonated regenerating the Zn-bound hydroxyl (Equation 2). This second step is regulated by the proton shuttle histidine (His) residue, which can adopt either an “in’ and an “out” conformation.

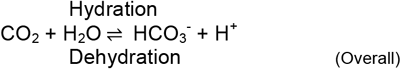

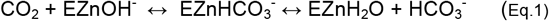

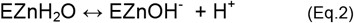

The α-class of CAs is made up of twelve catalytically active isoforms that differ in tissue distribution and subcellular localization (23). Of the human isoforms, only membrane bound carbonic anhydrase IX (CA IX) and carbonic anhydrase XII (CA XII) have been linked to cancer (10, 23) Both isoforms display their catalytic domains from the exofacial side of the plasma membrane (22). More attention has been placed on CA IX, as this isoform is upregulated by hypoxia, is associated with more aggressive forms of cancer, and is predictive of poor patient outcome (24–27). It also is considered a therapeutic target for many hypoxic tumors (28). There is additional evidence that CA IX forms functional interactions with specific ion transporters accelerating their activity (24, 29, 30). It has been hypothesized that CA IX forms a metabolon with these transporters and CA II to maintain the glycolytic phenotype by ‘cycling’ substrates between tumors cells and the microenvironment to regulated pH_i_ (10, 16, 23, 24, 29–31). Our lab, and others, have also hypothesized that CA IX serves to regulate pH_e_ by responding to changes in substrate concentration (10, 19, 20). As pH_e_ decreases, CA IX sequesters protons in the form of CO_2_ and H_2_O through the dehydration reaction (27, 32, 33). As pH_e_ increases CA IX produces HCO_3_^−^ and H^+^ via the hydration reaction (27, 33). Overall, this adjusts the pH to ~6.8. Exciting new data in the tumor setting supports this hypothesis, leading to the idea that CA IX serves as a pH-stat within the tumor microenvironment providing a survival and metastatic advantage to cancer cells that adopt the glycolytic phenotype (34). This role also prevents overacidification, which would ultimately lead to the death of both normal and cancer cells (34).

Previous work has shown that the MCT transporters interact with CA IX (24, 30). Recently it has been shown that expression of CA IX increases MCT transport activity by a non-catalytic mechanism involving the H^+^ shuttle mechanism mediated by His200 in CA IX (24, 30), which is analogous to His64 in the ubiquitous, cytosolic CA II isoform (35, 36). Theoretically, this non-catalytic mechanism would operate only when MCT’s are co-expressed with CA IX. We wondered if expression patterns of proton transporters, across cancer cells, could drive the ability of CA IX (or CA XII) to enhance proton efflux. To test this hypothesis, we first determined the expression of common ion transporters and the effect of hypoxia in cell models of breast cancer, including both triple negative (TNBC) and ER-positive luminal phenotypes. Our results show differential expression between and within different molecular subtypes of breast cancer cells most of which were insensitive to hypoxia. However, the TNBC line (UFH-001) that expresses exclusively CA IX, also express the MCT transporters both *in vitro* and *in vivo*. The same is true for a PDX model for TNBC. Knockdown of neither CA IX nor CA XII in UFH-001 or T47D (the ER-positive, luminal line that exclusively expresses CA XII) influenced ion transporter expression. Proton efflux measured using the glycolytic rate assay showed that UFH-001 cells were significantly more efficient in extruding protons than either control cells (MCF10A) or T47D cells. In addition, CA IX expression in UFH-001 cells and extracellular pH, independently influenced proton excretion rates. The same was not true for CA XII expression in T47D cells. These data support a role for CA IX, but not CA XII, in proton export. That said, sulfonamide-based inhibitors of CA IX did not affect proton efflux suggesting that CA IX regulation is via a non-catalytic mechanism. The coordinate expression of MCT4 and CA IX in aggressive breast cancers suggests a mechanism by which the proton shuttle function of CA IX may pass off protons to the MCT4 carrier to accelerate proton efflux.

## Experimental Section

### Cell culture and hypoxia

The MCF10A cells were plated at a density of 20,000 cells/mL in 10 cm dishes, and cultivated in in DMEM/Ham’s F12 medium, supplemented with 5% horse serum (Sigma Aldrich), 10 μg/mL insulin, 20 ng/mL epidermal growth factor (EGF) (Upstate Biochem) and 100 ng/mL dexamethasone (Sigma). T47D cells were plated in 10 cm dishes at 20,000 cells/mL and maintained in McCoys 5A medium supplemented with 10% FBS and 10 μg/mL insulin. MCF7 cells, plated in 10 cm dishes at 20,000 cells/mL, were maintained in DMEM supplemented with 10% FBS and 0.01 μM estrogen (Sigma Aldrich). UFH-001 and MDA-MB-231-LM2 cells were plated in 10 cm dishes at a density of 10,000 cells/mL and maintained in DMEM supplemented with 10% FBS. All cell lines were maintained at 37 °C in humidified air with 5% CO_2_.

Experiments were conducted when cells achieved ~70% confluency, unless otherwise specified. For hypoxia treatment, cells were placed in humidified Billups Rothenberg Metabolic Chambers and exposed to 1% O_2_, 5% CO_2_ and balanced N_2_ for designated times 37 °C. Desferroxamine mesylate (DFO) is an iron chelator that mimics hypoxia. For DFO treatment, a 10 mM stock was prepared in dH_2_O and filter sterilized. A final concentration of 100 μM DFO was used. All cell lines were authenticated and validated.

### RNA sequencing analysis

UFH-001 cells were exposed to either normoxic or hypoxic conditions (16 h). RNA isolated from these cells was checked for integrity using the Agilent Bioanalyzer 2100 system (Agilent Technologies, Santa Clara, CA) and quantified using a Qubit fluorometric assay (Thermo Fisher Scientific, Waltham, MA). Poly-A enriched mRNASeq libraries were prepared following Illumina’s TruSeq Stranded mRNA LT library preparation protocol (Illumina Inc., San Diego, CA) using 1 μg of total RNA. All samples were individually barcoded and quantitated with the KAPA Library Quantitation Kit for Illumina Platforms (Kapa Biosystems, Wilmington, MA) in conjunction with an Agilent Bioanalyzer DNA 1000 analysis (Agilent Technologies, Santa Clara, CA) for fragment size determination. The average fragment size was approximately 300 bp. 1.8 pM of the pooled libraries with 1% PhiX spike-in was loaded on one NextSeq 500/550 75 cycle High Output Kit v2 sequencing flow cell and sequenced on the Illumina NextSeq 500 sequencer. Filtered reads were mapped onto the Homo sapiens genome (Homo_sapiens.GRCh38.82) using Tophat2 v2.0.13, and processed by Cufflinks (v.2.2.1) for transcript abundance estimation and pairwise differential expression gene (DEG) analysis (37, 38). A total of 1,487 genes were upregulated and 1,459 genes were down regulated (p-value ≥ 0.05, Fold Change (FC) ≥ 1 and log_2_FC ≥ 0). A heat map was generated from a list of 623 DEG that had a log2 fold change > 1 and FDR (q-value) ≤ 0.05 using heatmap.2 in R. Individual FPKM (Fragments Per Kilobase per Million reads) expression values were used to investigate the transcriptional effect of hypoxia on key transporters and CA genes. An FDR (q-values) ≤ 0.05 was considered significant. The data discussed in this publication have been deposited in NCBI’s Gene Expression Omnibus (Edgar et al., 2002 (39)) and are accessible through GEO Series accession number GSE125511 (https://www.ncbi.nih.gov/geo/query/acc.cgi?acc=GSE125511).

### Xenograft and tumor graft models

All procedures were conducted in accordance with the NIH regulations and approved by the University of Florida IACUC. Both xenografts and a PDX model were used for the *in vivo* identification of proton transporters. For xenografts, a total of 300,000 UFH-001-Luc cells were suspended in culture medium and injected into the 4^th^ mammary gland of NOD/SCID mice (Jackson Laboratory), aged around 10-12 weeks old. The TNBC-derived tumor grafts were generated from cryo-preserved tissue (HCl-001), originally developed in Alana Welm’s lab (40), implanted into the 4^th^ mammary gland of NOD/SCID mice. When tumor volume reached 500 mm^3^, the mice were sacrificed, and dissected tumor was homogenized in RIPA buffer, containing protease inhibitor (Sigma Aldrich), using an Omni homogenizer. Protein concentrations were determined by the Markwell modification of the Lowry method and used to determine protein loading for SDS-PAGE analysis (41).

### Preparation of cell lysates and western blot analysis

Cells were washed 3x with ice-cold phosphate buffered saline [PBS, 120 mM NaCl, 2.7 mM KCl, 10 mM NaH_2_PO_4_.H_2_O and 10mM Na_2_HPO_4_ (pH 7.4)] and then exposed to RIPA buffer supplemented with protease inhibitor for 15 min on ice. Cell lysates were then scraped from the plates and clarified by centrifugation at 55,000 rpm for 60 min at 4°C. The clarified supernatants were collected and stored for protein analysis. Proteins were separated by SDS-PAGE and visualized using immunoblotting. Images were scanned and cropped (Adobe Photoshop version 11) for proper illustration of results. Antibodies were from the following sources: vATPase (Abcam, Ab103680), NHE1 (Santa Cruz, sc-58635), MCT1 (Santa Cruz, sc-365501), MCT4 (Santa Cruz, sc-50329), CAIX (M75 monoclonal, gift from Dr. Egbert Oosterwijk), CAII (Novus, NB600-919), CAXII (R&D, AF2190), and GAPDH (Cell Signaling, 5174). The GLUT1 antibody was made in the lab of SC Frost and has been previously characterized for specificity (42).

### Knockdown and deletion of CA expression

LentiCRISPR v2 was used to knockout CA IX expression in UFH-001 cells as previously described (27). GFP expression was monitored to confirm efficiency of transduction. Stably transduced UFH-001 and T47D cells were established by puromycin (2 μg/mL) selection (Sigma Aldrich). CA IX knockout and CA XII knockdown were confirmed by western blotting.

### Glucose transport assay

Glucose transport activity was measured according to previously published methods (43). For these experiments, we plated MCF10A and T47D cells, along with CA IX KO (UFH-001) cells and CA XII KO (T47D) cells at a density of 30,000 cells/35 mm plate. UFH-001 cells were plated at 10,000 cells/plate. At 50-75% confluency, the cells were divided into two groups. The control group was incubated under normal culture conditions, while the second group was exposed to 16 h of hypoxia. Cells were equilibrated in KRP buffer for 10 min at 37°C. Cells were then exposed or not to 40 μM cytochalasin B (a glucose transporter specific inhibitor that provides non-specific uptake), from a stock dissolved in DMSO for 10 min. This was followed by 10 min incubation with 0.2 mM [^3^H]-deoxyglucose. The reaction was terminated when cells were washed with ice-cold PBS and air dried at room temperature. To quantify radioactivity, the cells were lysed in 0.1% SDS and counted by scintillation spectrometry. Rates are presented as the concentration of [^3^H]-deoxyglucose uptake per min per plate.

### Glycolytic rate assay

MCF10A (5,000 cells/well), UFH-001 (2,500 cells/well) and T47D (10,000 cells/well) as well as the respective, CA IX KO (4,500 cells/plate) and CA XII KO (10,000 cells/plate) cells were seeded in 96 well plates and allowed to grow for 3 days. On day 3, medium was changed, and cells exposed to 16 h of hypoxia. On the day of the experiment, now day 4, cell culture medium was replaced with the glycolytic rate assay medium (XF Base Medium without phenol red plus 2 mM glutamine, 10 mM glucose, 1 mM pyruvate and 5 mM HEPES). For hypoxic cells, 100 μM DFO was added to the glycolytic rate assay medium. This protects HIF1a from degradation. These cells were then incubated for 1 h in the absence of CO_2_ and transferred to the Seahorse XF96 extracellular flux analyzer (Seahorse Bioscience). If present, ureido substituted benzene sulfonamide inhibitors (USBs) (5 μM), dissolved in assay medium, were administered via the first injection port for acute treatment. A rotenone/antimycin cocktail (Rot/AA) and deoxyglucose (DG) were placed in the second and third ports, respectively, and added sequentially during the assay. Glycolytic rate was measured according to the manufacturer’s instructions and results analyzed using the XF report generator.

### Carbonic anhydrase inhibitors

CA catalytic activity was blocked with three USB compounds; U-CH_3_ (4-**{[**3,5-methylphenyl)carbamoylamino}benzenesulfonamide), SLC-0111/U-F (4-**{[**4-fluorophenyl)carbamoyl]amino}benzenesulfonamide), and U-NO_2_ (4-**{[**3-nitrophenyl) carbamoyl] amino}benzenesulfonamide). The synthesis and K_i_ values for recombinant CA proteins are described elsewhere (44). K_i_ values for inhibitors in intact breast cancer cells have also been previously reported (45). 100 mM stock concentrations of these inhibitors were prepared in DMSO and diluted to specific concentrations as specified in the Results section. The final DMSO concentration was ≤ 0.5%.

### Lactate determination

MCF10A and T47D cells, along with CA IX KO (UFH-001) cells and CA XII KO (T47D) cells were plated at a density of 30,000 cells/35 mm plate, while UFH-001 cells were plated at 10,000 cells/plate. All cells were grown to 50-70% confluence. Cells were fed with appropriate medium and then exposed to 16 h of hypoxia at different pH values. Medium was collected from each plate at the end of the exposure time and immediately assayed for lactate. Alternatively, cells were handled similarly as for the glycolytic assay, i.e., fresh medium, adjusted to either pH 7.4 or 6.8, was provided followed by a 3 h incubation at 37°C in a CO_2_ incubator at which point medium samples were collected for lactate analysis (46). For those cells initially exposed to hypoxia, DFO was added as a chemical substitute. Lactate was determined using a coupled enzyme reaction. Data are reported as nmol normalized to protein concentration.

### Structural representation of the interaction between USB compounds and CAs

X-ray crystallography structures of USB compounds in complex with CA II and a CA IX mimic (analogous site generated mutagenesis of residues in the active site of CA II to resemble wildtype CA IX) were previously obtained and the data deposited in the protein data bank (35). *In silico* modelling experiments of the same USBs modeled into the active site of CA XII were also previously performed and published (45). These structures were used to illustrate USB binding in the active sites of CA II, IX and XII relative to His64 (CA II numbering). Figures were made using PyMol.

### Statistics

RNA seq data for normoxic vs hypoxic cells, along with the CA activity assay, were analyzed using the Student’s t test. All *p* values were based on two-tailed analysis and *p* < 0.05 was considered statistically significant. Statistical analysis was performed using the Prism 7 software. Glucose transport activity, glycolytic activity (GlycoPER), extracellular lactate, and pH_e_ data were analyzed with R (version 3.4.1) using RStudio (V.1.0.153). ANOVA models were generated with the lm function using type III Sum of Squares, and Tukey posthoc comparisons among groups were conducted with the LSD test function of Agricolae.

## Results

### The MCT proton transporter family is coordinately expressed with CA IX in breast cancer cells and tissue

To compare the expression profiles of ion transporters with molecular subtypes of breast cancer, we compared protein expression of a specific set of transporters, including V-ATPase, NHE1, and MCT1/4, in selected breast cells lines and their correlation with CA IX and CA XII expression (Figure 1A). These data show that transporter expression varied across cell lines and under different conditions. We examined two TNBC cell lines: UFH-001 and MDA-MB-231-LM2. In UFH-001 cells, but not MDA-MB-231 cells, there was strong MCT1 and MCT4 expression in cells maintained under normoxic conditions (N) with limited sensitivity to hypoxia (H) (Figure 1A). In our hands, only the UFH-001 cells expressed CA IX, which was significantly upregulated by hypoxia as we have shown before (26, 27) and was the only cell line that expressed CA II. That the MDA-MB-231-LM2 cells did not express CA IX was surprising based on previous findings with MDA-MB-231 cells (47). The LM2 cells represent a line that was developed from a metastatic site in the lungs that derived from an MDA-MB-231 tumor generated in mice (48) but has not been previously tested for CA IX expression. We also examined two luminal ER-expressing cell lines: T47D and MCF7. The T47D cells displayed strong V-ATPase and NHE1 protein expression under normoxic conditions. Neither transporter showed sensitivity to hypoxia. These cells showed exclusive expression of CA XII, which was not induced by hypoxia (Figure 1A). In contrast, the MCF7 cells showed low V-ATPase and MCT1 expression, low expression of CA XII, but inducible CA IX expression (Figure 1A). This later feature, hypoxia-dependent CA IX expression, excluded them from further analysis because of the difficulty in interpreting data in cells with multiple cell surface CAs. The MCF10A cells, used as a control in our studies, showed hypoxia-dependent upregulation of NHE1, but no other proton transporter. They did exhibit hypoxic-dependent expression of CA IX protein in the absence of CA XII (Figure 1A). Although not shown here, we have previously published data that demonstrates that the breast cancer cells that are the specific focus of this study show hypoxic-dependent HIF1α protein expression (43)

**Figure 1.**
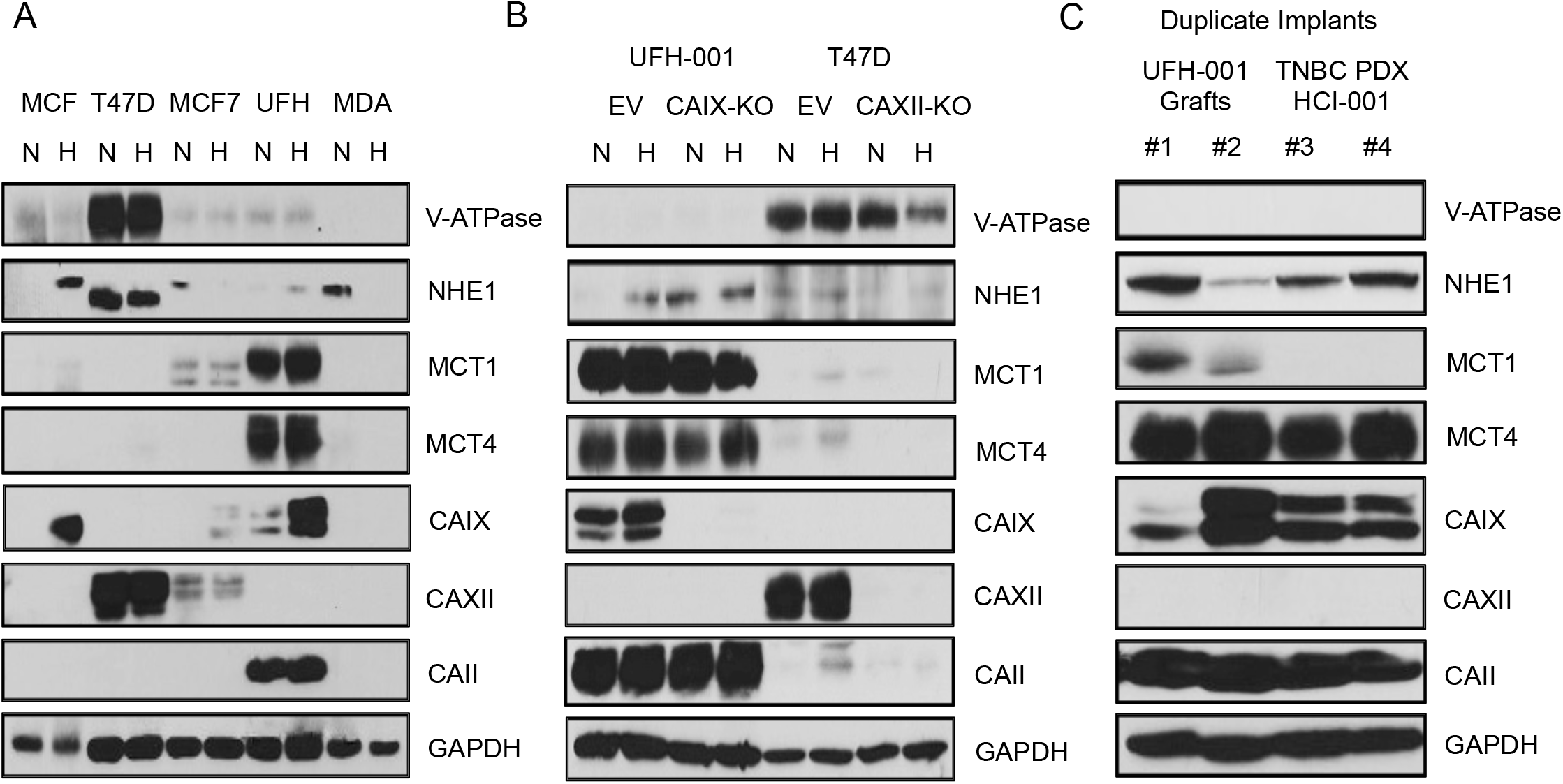
Cell specific protein expression of transporters in TNBC and luminal breast cancer cell lines and tumor grafts. Panel A) Breast cell lines were grown to ~75% confluence and then exposed to normoxic (N) or hypoxic conditions (H, 1% O_2_ for 16 h). After 16 h, the cells were rinsed with PBS and lysates extracted using RIPA buffer containing protease inhibitors. Equal concentrations of protein were loaded onto SDS PAGE gels, and then transferred to nitrocellulose for western blot analysis. Panel B) Frozen tissue samples from UFH-001 derived xenografts (#1 and #2) and patient derived tumorgrafts (#3 and #4) were homogenized in RIPA buffer containing proteinase inhibitor, resolved on a 10% SDS-PAGE gel and then transferred to nitrocellulose for western blot analysis. Panel C) UFH-001 and T47D empty vector (EV) controls as well as their respective CA IX and CA XII KO cells were cultured and then exposed to either N or H (16 h). After which cell lysates were collected and equal volumes of proteins loaded onto SDS PAGE gels for western blot analysis. Membranes were probed for CA expression (CA IX, CA XII and CA II), and transporter expression (V-ATPase, NHE1, MCT1, MCT4 and GLUT-1). GAPDH was used as a loading control.

To determine if transporter expression was affected by the presence or absence of membrane bound CAs, we utilized RNAi strategies to knockdown CA IX expression in the UFH-001 cells and CA XII in the T47D cells (Figure 1B). We selected these cells because of their specific expression of CA IX and CA XII, respectively. When compared to empty vector (EV) controls, our data showed that knockdown of CA IX or CA XII expression had little effect on the expression of proton transporters.

To evaluate the potential for proton transporter function, *in vivo*, we generated xenografts of UFH-001 (duplicate tumors #1 and #2, Figure 1C) and TNBC patient derived tumor grafts (PDX) (duplicate tumors, #3 and #4, Figure 1C). In UFH-001 xenografts, both isoforms of MCT were expressed, although MCT4 expression patterns were higher relative to MCT 1. NHE1 expression was variable but appeared more highly expressed than in the cultured cells. There was a total lack of V-ATPase which is consistent with the data in UFH-001 cell culture. CA IX expression was somewhat variable in the tumors derived from UFH-001 cells, perhaps because of the extent of local hypoxia. CA II expression was strong for both UFH-001 xenograft and PDX models. In the TNBC PDX model, there was strong MCT4 expression, but essentially no MCT1. NHE1 was present but again V-ATPase was not observed. CA IX expression was consistently high. No expression of CA XII was detected in either the UFH-001 xenografts or the TNBC PDX model. We had difficulty growing either the T47D xenografts or a PDX model for luminal breast cancer, so we are unable to make comments about the expression of proton transporters (or CA XII expression) in *in vivo* luminal models. What we can conclude is that the expression pattern of both ion transporters and CA IX expression in the tumor grafted UFH-001 cells and the TNBC PDX model are consistent with the observations in cultured UFH-001 cells.

An RNA sequencing experiment using the UFH-001 cells under normoxic and hypoxic conditions support the protein expression pattern. A total of 1,487 genes were upregulated under hypoxic conditions and 1,459 were down regulated (p-value ≤ 0.05). A heat map was generated from 623 DEG (differentially expressed genes) that had log2 scores > 1 and FDR (false discovery rate, or q-value) ≤ 0.05 (Figure 2A). This showed a distinct transcriptional profile between normoxic and hypoxic conditions. From these data bases, we selected information specific to the ion transporters and CA family members (Figure 2B). Very few transcripts of NHE1 and V-ATPase were detected and expression of neither was sensitive to hypoxia (Figure 2B). MCT1 and MCT4 showed higher level of transcripts than those for NHE1 and vATPase. MCT1 transcription was significantly upregulated by hypoxia (q-value = 0.0013) but this did not translate into higher levels of MCT1 protein (Figure 1A). Also, of note is the strong transcription of CA IX and CA II. Hypoxia increased CA IX transcription by over 5-fold (q-value = 0.0013), while CA II transcription decreased but not significantly (q-value = 0.0817) (Figure 2B). While CA XII gene expression was extremely low (Figure 2B), hypoxia increased CA XII transcription by nearly 3-fold (q value = 0.0013). Despite this, we never observe CA XII expression at the protein level in UFH-001 cells (or UFH-001 xenografts or the TNBC PDX model).

**Figure 2.**
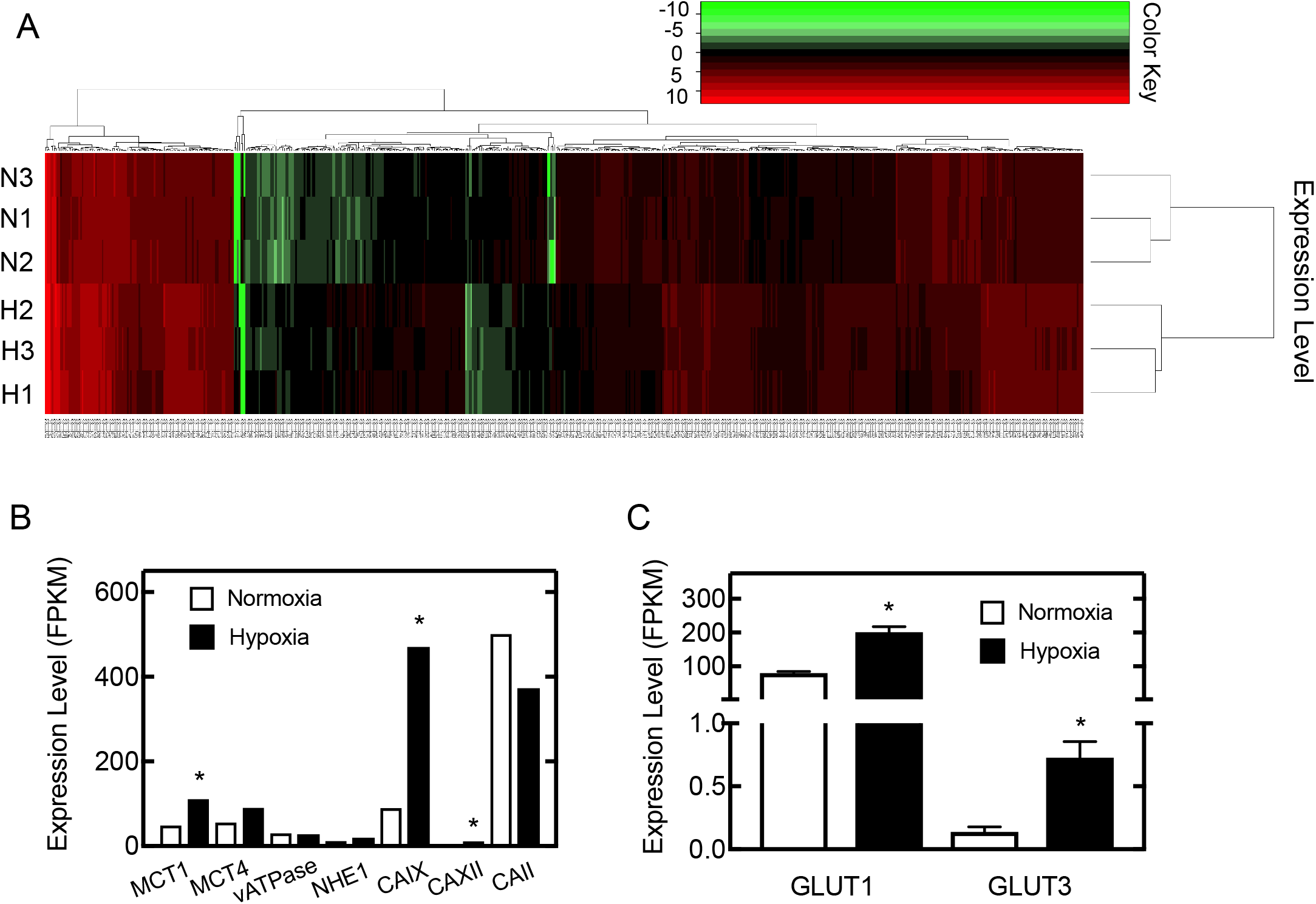
RNA sequencing support protein expression data. Panel A) A heat map was generated from a list of 623 differentially expressed genes (DEG) that had a log2 fold change > 1 and FDR (q-value) ≤ 0.05. Color key is shown separately. Red represents upregulated genes; green represents downregulated genes. Panel B) FPKM values are represented for the ion transporters of interest, along with three members of the carbonic anhydrase family. Panel C) FPKM values for two of the glucose transporter family: GLUT1 and GLUT3. * represents q ≤ 0.05 for hypoxia vs normoxia.

### Proton transporter expression is associated with survival in breast cancer patients

Breast cancer patient survival is often predicted by gene expression patterns. To determine the association of specific members of ion transporter family with patient outcome, we examined publicly available microarray data through the Kaplan-Meier database (kmplot.com/analysis/). We used the web tool developed by Lanczky *et al*. to analyze these databases (49). Supplemental Figure 2A shows that overexpression of V-ATPase is a positive prognosticator when queried across all breast cancer subtypes as a group. Overexpression of NHE1 also predicts better outcomes (Supplemental Figure 2B). In contrast, MCT1 overexpression is associated with decreased survival (Supplemental Figure 2C) and MCT4 is an even stronger predictor than MCT1 of poor patient survival (Supplemental Figure 2D). This suggests that the MCT transporter family is associated with more aggressive types of cancer and is associated with CA IX expression (Figure 1), another marker for poor patient outcome.

### CA IX expression, but not that of CA XII, regulates proton efflux in UFH-001 cells

The glycolytic rate assay (developed by Seahorse Biosciences) was used to determine proton excretion rates from glycolysis (GlycoPER) in UFH-001 and T47D cells relative to the control MCF10A cells (Figure 3). This assay simultaneously measures extracellular acidification rates (ECAR) and oxygen consumption rates (OCR). In this version of the assay, cells are provided glucose (10 mM) at the start of the experiment (thus the high initial rates of ECAR in Figure 3A) and then treated with a cocktail containing rotenone/antimycin (Rot/AA) to block oxidative phosphorylation followed by deoxyglucose (DG) to block glycolysis. Raw data are shown in Figures 3A and 3B for these two parameters. Figures 3C and 3D represent the calculated rates of proton extrusion that subtract out the contribution of CO_2_ derived from mitochondrial activity to extracellular acidification. The resulting value, GlycoPER, is the rate of protons extruded into the extracellular medium presumably from glycolysis in real time. To maintain the hypoxic phenotype in the absence of a hypoxic environment, the hypoxic state during the assay was mimicked by the addition of 100 μM desferroxamine mesylate (DFO) to maintain the stability of HIF1α. CA and ion transporter expression, in the presence of DFO, were comparable to those observed under hypoxic conditions (Supplemental Figure 2). We also exposed cells, during the assay, to buffer that was initially at pH 7.4 or pH 6.8, the latter of which is observed in hypoxic tumors. We have shown previously that these different pH conditions alter the activity of CA IX and CA XII, which might change the rate of extracellular proton production based on changes in CA activity (which we confirmed here: see Figure 5).

**Figure 3.**
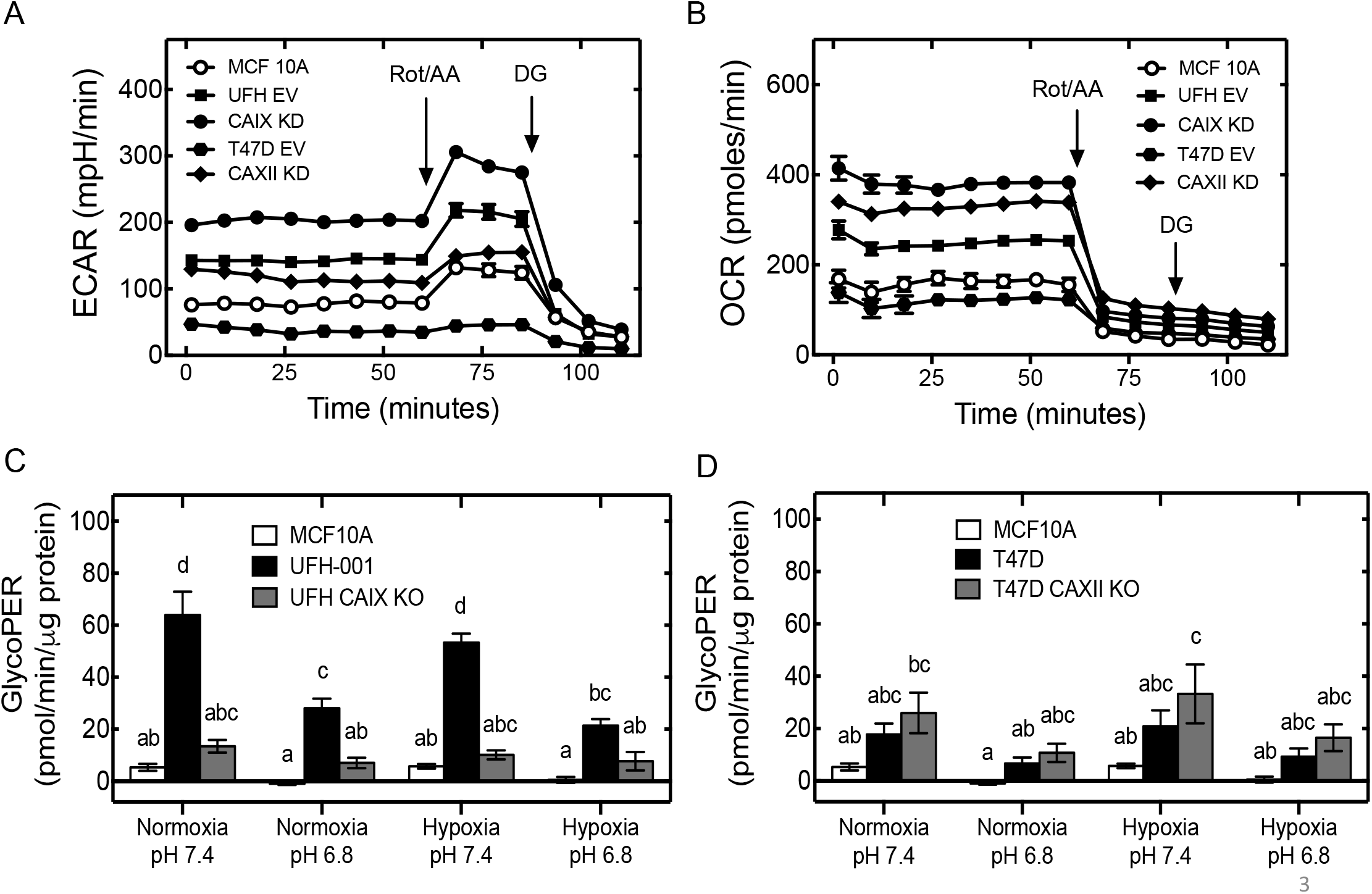
The glycolytic phenotype in breast cancer cells. Panel A) Extracellular acidification rate, ECAR and Panel B) oxygen consumption rate (OCR) were determined in MCF10A, UFH-1, T47D and the respective CA IX and CA XII KO cells after sequential compound injections: Rot/AA (rotenone and antimycin cocktail) and DG (deoxyglucose). Proton transport rates from glycolysis (GlycoPER) was assessed in Panel C) UFH-001 empty vector control (EV) and CA IX KO versus MCF10A cells and in Panel D) T47D empty vector control (EV) and CA XII KO cells versus MCF10A cells, under different pH conditions having been exposed to either normoxic or hypoxic conditions for 16 h prior to the Seahorse assay. n = 3 for all data sets.

GlycoPER data for UFH-001 cells and T47D cells are shown in Figures 3C and 3D respectively. In each plot, data are compared to the MCF10A cells, under all tested conditions. Under normoxic conditions at pH 7.4, UFH-001 cells produce significantly higher proton efflux rates relative to MCF10A cells (Figure 3C). The same is observed under hypoxic conditions. At pH 6.8, the efflux rate in UFH-001 cells is significantly reduced relative to pH 7.4 under both normoxic and hypoxic conditions. CA IX knockout further reduced GlycoPER relative to EV controls under all conditions. We conclude that pH and CA IX expression have independent effects on GlycoPER. The GlycoPER pattern in T47D cells was strikingly different from UFH-001 cells (Figure 3D). In fact, there was little statistical significance across the conditions, although an upward trend in proton efflux was observed in CA XII knockdown cells relative to EV control T47D cells. The GlycoPER rate in UFH-001 EV cells was about 2-3 times higher than that in T47D cells. Overall, these data suggest that CA IX expression, but not that of CA XII, contributes to extracellular acidification

### CA expression does not influence glucose transporter expression or activity

Cells in culture exhibit increased glycolysis perhaps because they have an unrestricted source of glucose. Cancer cells also exhibit increased glucose uptake and enhanced glycolytic rates as part of the glycolytic switch (4). This is due, in part, by increased constitutive expression of GLUT1, which consequently increases glucose uptake into the cell (50). We demonstrate in Figure 2C that the level of GLUT1 transcripts in UFH-001 cells was high and that the number of transcripts increased in response to hypoxia (~ 2-fold). GLUT3, although transcript number was lower by nearly 1000-fold relative to GLUT1, was strongly regulated by hypoxia. It seems unlikely that GLUT3 contributes significantly to glucose uptake in these cells, although we did not analyze GLUT3 protein expression. We have confirmed the strong protein expression of GLUT1 in UFH-001 and the control MCF10A but much lower expression in the luminal T47D cells (Figure 4A). We also show that GLUT1 expression in both the UFH-001 and MCF10A cells was sensitive to hypoxia. Neither knockdown of CA IX in UFH-001 cells nor CA XII in T47D cells affected GLUT1 expression. To analyze glucose transport in these cells, we measured uptake of [^3^H]2-deoxyglucose which is trapped as [^3^H]2-deoxyglucose-6 phosphate in the cells. Figure 4B demonstrates that basal glucose transport in UFH-001 cells was greater than that observed for either MCF10A or T47D cells. Hypoxia increased uptake but for the most part, these values were not statistically significant. We have observed this trend before (43) which may reflect the high constitutive expression of GLUT1 induced by the transformation process itself. Knockdown of CA IX or CA XII in UFH-001 and T47D cells, respectively, did not significantly alter glucose uptake relative to EV controls. Therefore, the effect exerted by CA IX on proton efflux is independent of glucose.

**Figure 4.**
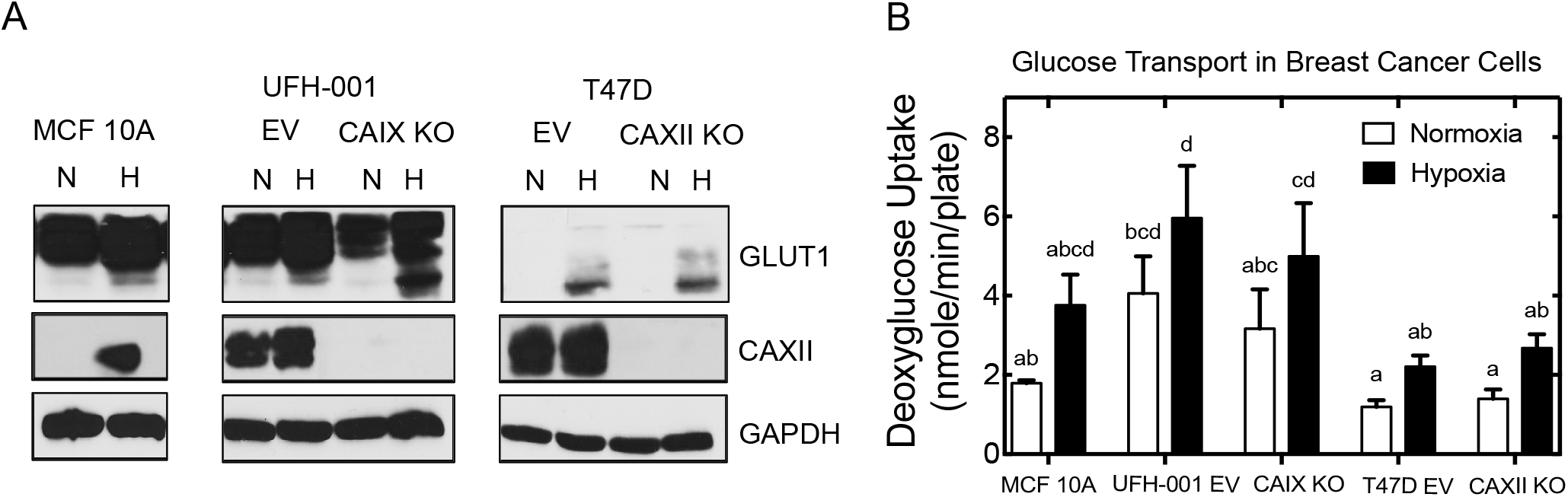
GLUT1 expression and transport activity in breast cancer cells. Panel A) CA and GLUT1 expression were probed for in lysates isolated from breast cancer cell lines under normoxic conditions or 16 h of hypoxia. N = normoxia, H = hypoxia. GAPDH was used as a loading control. Data are representative of 3 independent experiments. B) Deoxyglucose uptake was assessed in breast cell lines treated as described in Panel A. n = 3.

### Low pH increases activity of both CA IX and CA XII

To determine the effect of pH on CA activity, we have calculated the first order rate constants for CA IX activity in UFH-001 cells (Figure 5A) and CA XII activity in T47D cells (Figure 5B) using membrane inlet mass spectrometry (MIMS). Low pH (6.8) significantly increased the activity of CA IX and CA XII relative to pH 7.4 in their respective cells. The fold increase in activity in UFH-001 cells was ~ 2.5-fold in normoxic cells and almost 4-fold in cells exposed to hypoxia. The fold increase in CA XII activity in T47D cells was considerably less than changes in CA IX activity: about 50% in both normoxic and hypoxic cells. In previous studies, we have shown that the increase in activity at low pH is related in part to the acceleration of the dehydration activity of CA IX (favoring proton consumption) relative to its hydration activity (producing protons). That low pH is less influential with respect to CA XII may be related to its reduced stability compared to that of CA IX which is stable to values of pH as low as 3.0 (51).

**Figure 5.**
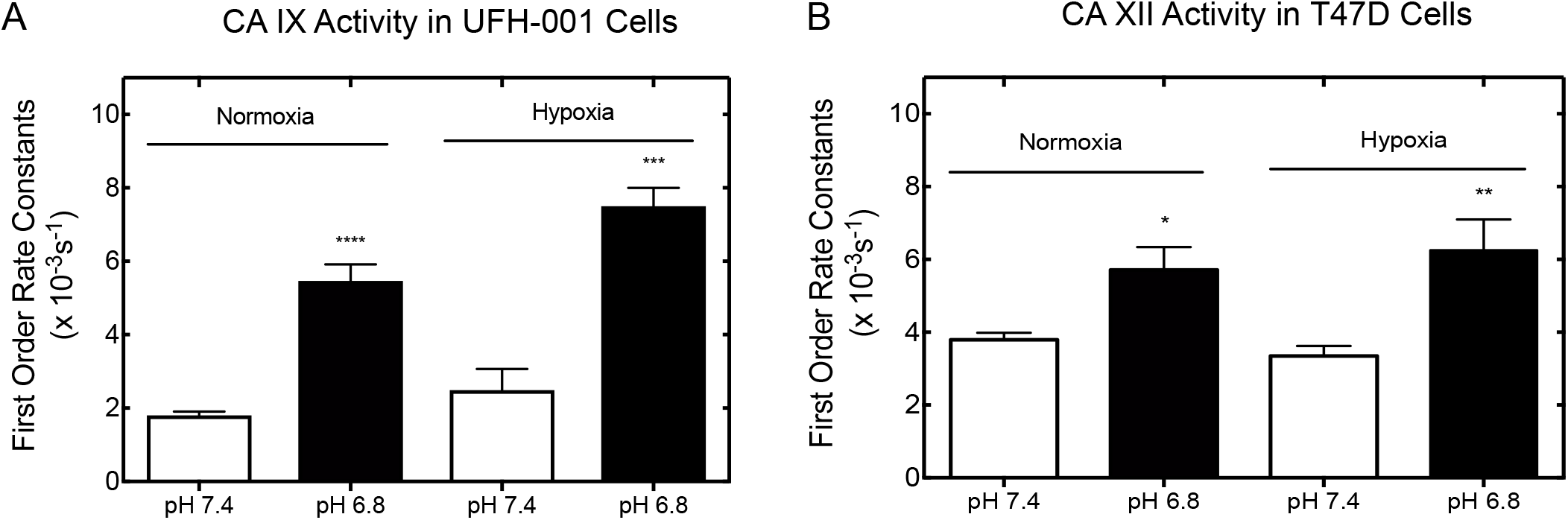
Alterations in CA IX and CA XII activity in response to pH and hypoxia. Panel A) CA IX activity in UFH-001 cells, under low (6.8) or physiological (7.4) pH and under normoxic or hypoxic conditions (16 h), was measured using the MIMS assay and first order rate constants calculated from these data. ****p < 0.0001, ***p=0.002 (pH 6.8 vs 7.4 for normoxic and hypoxic cells, respectively). Panel B) CA XII activity, under low (6.8) or physiological (7.4) pH and under normoxic or hypoxic conditions, was measured using the MIMS assay and first order rate constants calculated from these data. *p = 0.047, **p = 0.026 (pH 6.8 vs 7.4 for normoxic and hypoxic cells, respectively). n = 3 for all data sets.

### CA catalytic activity is not required for decreased proton efflux in breast cancer cells

Ureido substituted benzene sulfonamides (USBs) are potent inhibitors of CA IX and CA XII, over the off-target cytosolic and ubiquitously expressed CA II, as measured by stop flow kinetics (SFK) of purified recombinant proteins (35, 44). Using MIMS, we have shown that these same inhibitors selectivity target and inhibit CA IX activity over CA XII in UFH-001 cells versus T47D cells, respectively (45). Thus, we have used these inhibitors (the structure of which are shown in Figure 6A) to determine if the reduction of GlycoPER in CA IX KO cells, relative to the EV control UFH-001 cells, is related to its catalytic activity (Figure 6B). This does not appear to be the case, as the inhibitors did not significantly affect the rate of GlycoPER under either normoxic or hypoxic conditions. We conclude that the expression of CA IX, but not its activity, influences proton efflux/acidification.

**Figure 6.**
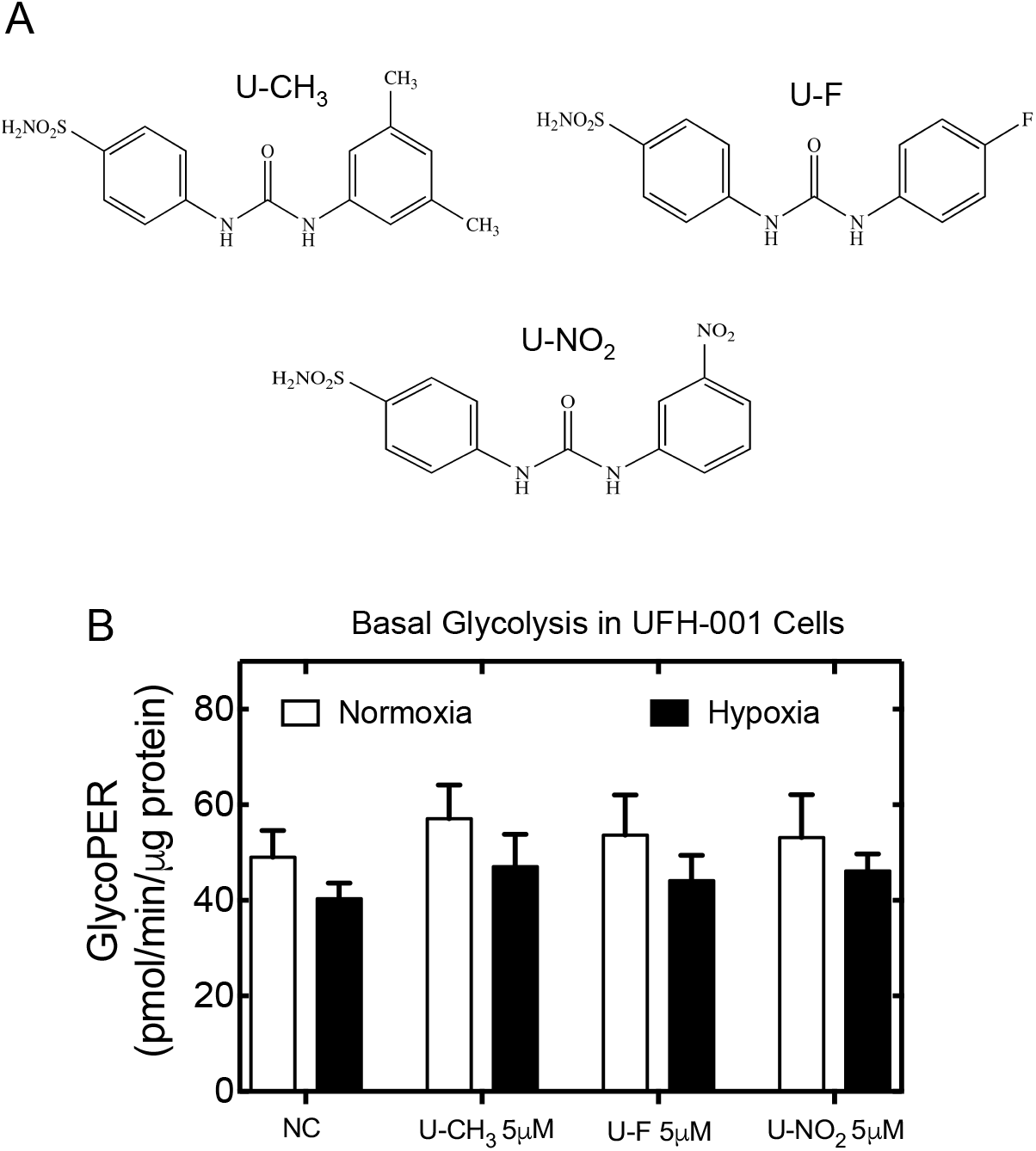
Effect of ureido sulfonamides (USBs) on GlycoPER in UFH-001 cells. Panel A) Structures of USBs and their abbreviations. Panel B) UFH-001 cells were exposed to normoxic or hypoxic conditions for 16 h. Cells were then prepared for the Seahorse assay and during that time exposed to inhibitors (5 μM) during the duration of the assay. This inhibitor concentration is well above the K_i_ values for the inhibition of CA IX and CA XII in intact cells. n = 3.

Glucose transport rates in UFH-001 cells were not affected by the USBs, either (Supplemental Figure 3A). The same was true for glucose transport activity in T47D cells (Supplemental Figure 3B). Again, these data suggest that neither CA IX nor CA XII directly affects glucose uptake.

### CA expression does not affect extracellular lactate concentrations in breast cancer cells

Glucose metabolism in cancer cells is accelerated and leads to the formation of lactic acid because of the switch from aerobic to anaerobic metabolism. This leads to reduced extracellular pH through export of lactate. Here, we determined if changes in pH or CA expression affect extracellular lactate concentrations (Figures 7A and 7B) under conditions that replicate the protocol used for the GlycoPER assays (Figures 3A and 3B). We first exposed cells to either normoxic or hypoxic conditions for 16 h. Then, cells were washed and exposed additionally to normoxic conditions or DFO for 3 h at different pH values. In Figure 7A, we show that lactate export in UFH-001 was significantly higher under all conditions relative to the MCF10A controls, similarly to the increase in proton efflux as might be expected (Figure 3C). However, there were some striking differences between the GlycoPER data and the lactate measurements in the UFH-001 cells with respect to pH, and the knockdown experiments. pH had no effect on lactate levels, and knockdown of CA IX trended toward an increase in lactate, although not significantly. Yet, both reduced pH and loss of CA IX independently reduced the apparent proton efflux in the GlycoPER assay (Figure 3C). Lactate concentrations in the medium from T47D cells were also significantly higher than that of MCF10A cells under all conditions (Figure 7B). However, neither hypoxia, pH, nor CA XII knockdown affected lactate concentrations. These data are similar to the proton efflux as presented in the GlycoPER results for T47D cells (Figure 2D). These data suggest that the effect of CA IX on proton efflux is independent of not only glucose uptake but glycolysis, as well.

**Figure 7.**
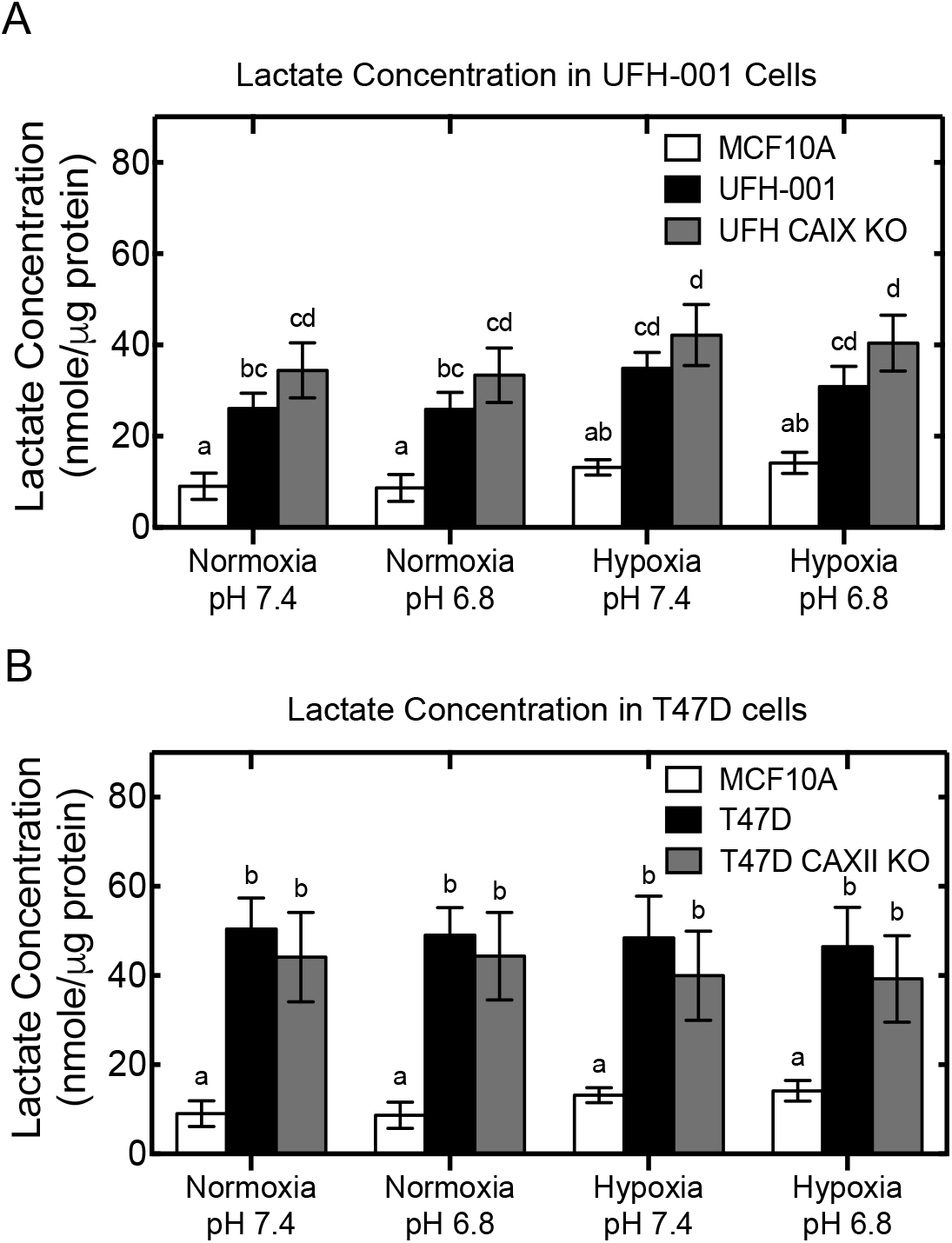
Lactate concentrations in breast cancer cells. Panel A) UFH-001 cells (EV and CA IX KO cells) and MCF10A cells were exposed to normoxic or hypoxic conditions for 16 h. Cells were then incubated for another 3 h at either pH 7.4 or pH 6.8, after which lactate concentrations was determined. During those 3 h, hypoxic cells were exposed to desferroxamine mesylate DFO (100 μM). Panel B) T47D cells (EV control and CA XII KO cells) and MCF10A cells were exposed to normoxic or hypoxic conditions for 16 h. Cells were then incubated for another 3 h at either pH 7.4 or pH 6.8, after which lactate concentrations was determined. During those 3 h, hypoxic cells were exposed to DFO (100 μM). n = 3 for all data sets.

## Discussion

Because proton export is dependent on transporters, our first goal was to determine the expression patterns of these across luminal and triple negative breast cancer cells. Our studies show that members of the ion transport family are differentially expressed in breast cancer cells (Figure 1). Of interest was the expression of the MCT4, lactate transporter (which favors lactate export), and its preferential expression in the aggressive, UFH-001 cells, and its apparent coexpression with CA IX. This combined expression was preserved in the UFH-001 xenografts and the PDX (TNBC) model, strengthening the relationship between these two proteins, both of which are poor prognosticators for patient outcome according to the Kaplan Meier database. Interestingly, Lee *et al*. have shown that increased lactate levels are associated, *in vivo*, with overexpression of CA IX (34), although our *in vitro* studies do not show a correlation between lactate levels and CA IX expression (Figure 7A). However, Lee *et al*. also showed that most of the increases in total lactate levels in the tumor setting resulted from intracellular lactate accumulation, with no difference in extracellular lactate concentrations between cells in which CA IX was overexpressed compared to controls (34). Perhaps these data explain the lack of difference in extracellular lactate levels between our CA IX expressing UFH-001 cells and those in which CA IX expression is ablated. The UFH-001 cells also express MCT1, but its expression is reduced in UFH-001 xenografts relative to MCT4 and is absent in the triple negative PDX model. This suggests that MCT4 may to the major lactate transporter, *in vivo* in TNBC. In contrast, it appears that the V-ATPase is associated with luminal breast cancer and CA XII expression. Based on the Kaplan Meier data from all breast cancer patients (Supplemental Figure 1), MCT4 expression is associated with poor patient outcome, while V-ATPase is associated with better prognosis, consistent with the differential expression pattern shown in our study (Figure 1). The T47D line expresses little MCT1 or MCT4 yet extracellular lactate concentration is surprisingly similar to that of the UFH-001 cells. Perhaps, these cells express one of the other MCT family members (12) or less lactate accumulates inside the cells.

The glycolytic rate assay revealed that CA IX expression is associated with high rates of proton export relative to either the control MCF10A cells or the T47D cells at pH 7.4 (Figure 3). Because MCT1 and MCT4 are the only ion transporters overexpressed in the UFH-001 cells, it is likely that they are responsible for both lactate and proton transport. Catalytically, CA IX at pH 7.4 might be expected to participate in extracellular proton production (19, 20, 27, 34, 45) as this pH favors the hydration reaction leading to the production of protons in the extracellular space (33). Indeed, it appears that proton production increases in normal UFH-001 cells, at pH 7.4, based on the GlycoPER assay (Figure 3C). We would also expect that at pH 6.8, where the dehydration reaction is favored (33), that proton production would decrease. This is exactly what we observed, in fact, under both normoxic and hypoxic conditions in the GlycoPER assay (Figure 3C). These data support the studies of Swietach *et al*. and Lee *et al*. who have shown that CA IX has a stabilizing effect, in that CA IX can participate in both acidification and pH control (a pH stat) of the tumor microenvironment to adjust the pH to one that favors cancer cell viability both *in vitro* and *in vivo* (19, 20, 34).

That said, blocking CA activity with USBs had no effect on proton efflux (Figure 6). Given this result, it was surprisingly that knockdown of CA IX reduced the rate of proton production in the extracellular space, under all conditions, to values that were not statistically different from the values observed at pH 6.8 (Figure 3C). This suggests that without CA IX protein proton flux is limited. This questions the role of CA IX catalytic activity in this setting. Jamali *et al*. (24) have demonstrated that CA IX augments MCT1 and MCT4 transport activity, but not expression, the latter of which is corroborated by our present studies. These authors also proposed that the effect of CA IX on MCT function occurs via a non-catalytic mechanism (24). Specificallly, they showed that the histidine involved in proton shuttling in CA IX (Equation 2) is important in collecting intracellular protons and in accelerating their export through MCT. In CA IX, this histidine is found at position 200 (24). The CA isoform that has been well studied, CA II, has this histidine at position 64 (36). We have used the CA II numbering system to show the position of this histidine in structural models of CA II (Panel A), CA IX (Panel B), and CA XII (Panel C) in Figure 8. It is clear that this histidine occupies a separate space (the hydrophilic side of the catalytic pocket) from the USB inhibitors (which bind in the hydrophobic pocket). This suggests that the proton transport function of this residue can act independently from the hydration/dehydration reaction (Equation 1). Thus, CA IX could sequester protons through this “antenna” and pass them on to the MCT transporters as suggested by the Becker group (24, 30) independent of its catalytic activity. CA IX also contains a unique proteoglycan (PG) domain, which is negatively charged. Ames *et al*. have shown that the PG domain also facilitates MCT transport activity (30). The PG domain is absent from CA XII which may underlie our observation that in the T47D cells, which express only CA XII, proton efflux is not influenced by CA XII knockdown, even though CA XII contains an analogous His64 (at position 94) (22). This suggests that both the antenna and the PG domain play important roles in proton-coupled lactate transport in cancer cells that express CA IX.

**Figure 8.**
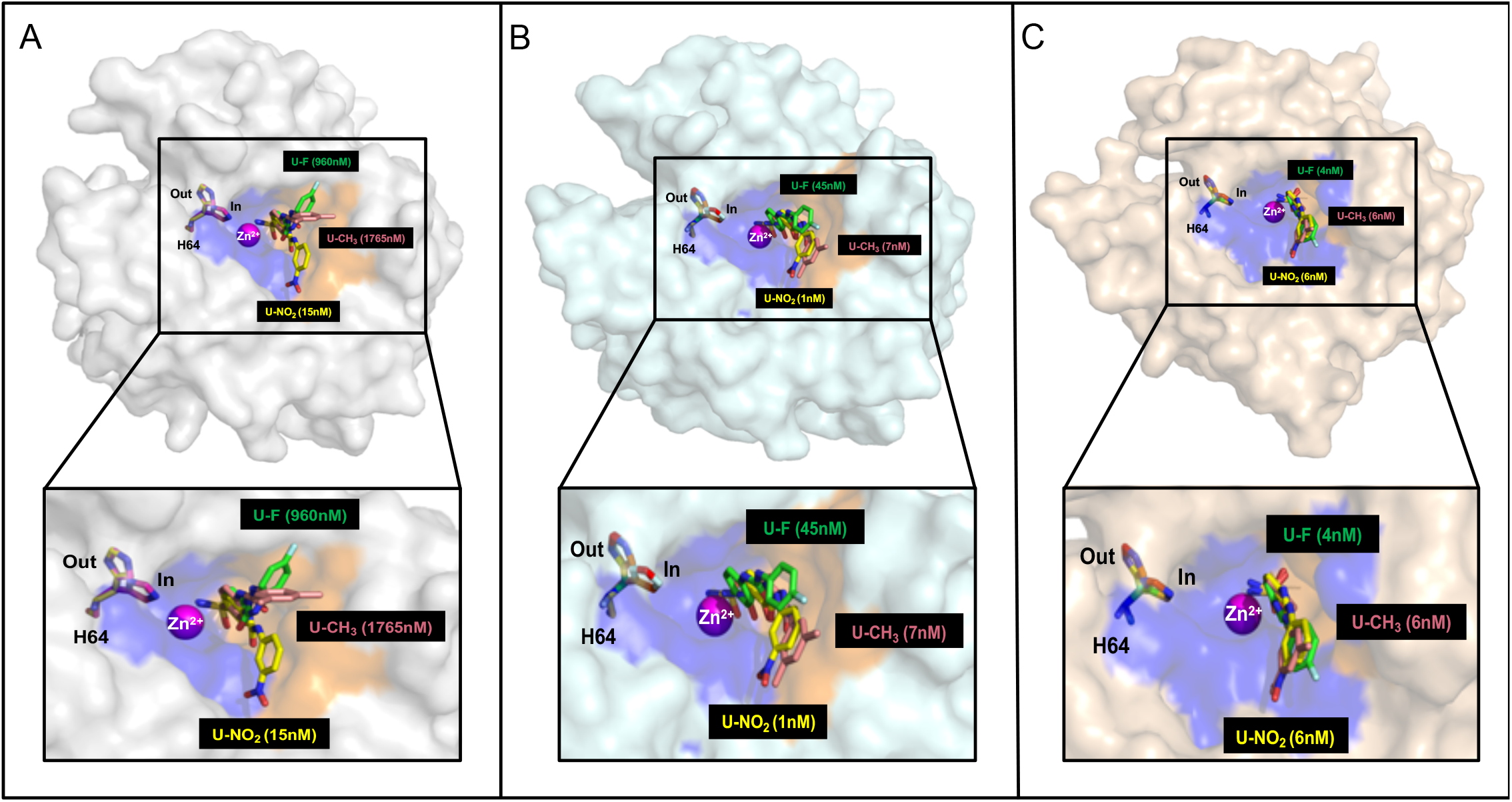
USBs bound in the active site of CAs. Surface representation of compounds U-CH_3_ (pink), U-F (green) and U-NO_2_ (yellow) in complex with monomers of CA II (gray), the CA IX-mimic (cyan), and modeled into the active site of CA XII (wheat). The catalytic zinc (magenta sphere) and His64 (H64, mutilcolored, in and out positions) are also shown. CA II numbering is used in these structural models. The analogous histidine in CA IX is at position 200, and at position 94 in CA XII.

Overall, the fact that CA IX knockdown, but not inhibition of CA IX catalytic activity, affects proton flux leads us to support the model that emphasizes the importance of the proton shuttle activity of His200 located in catalytic domain of CA IX and the PG domain, the unique extension of CA IX (52). Thus, His200 and/or the PG domain that may facilitate proton transfer to the monocarboxylate transporters (MCT1 and MCT4) to enhance proton-dependent lactate flux. A lot of attention has been placed on CA IX as a therapeutic target in hypoxic tumors (28), and many of the drugs are based on the interaction of sulfonamide-based inhibitors with Zn^++^ in the catalytic pocket. We speculate that CA IX-specific catalytic inhibitors should be re-designed to not only inhibit the catalytic function associated with the catalytic pocket, but also inhibit the proton shuttling function of His200. Alternatively, inhibitors against the MCT transporters could be used in combination with CA IX specific sulfonamide inhibitors to overcome the extracellular acidification that favors cancer cell growth and metastatic behavior.

## Acknowledgements

The authors would like to recognize the exceptional cell culture skills of Xiao Wei Gu. We would also like to thank the Center for Immunology and Transplantation at the University of Florida for access to the Seahorse equipment.

## Author contributions

MYM, ZC, DK, AW, and LA performed experiments. CDH provided oversight for xenograft and PDX animal experiments. MB, FC, and CTS provided sulfonamide inhibitors. KB created the Crispr CA IX knockout cells. CJF performed the statistical analysis and analyzed the RNA seq data. MYM, RM, and SCF developed the research strategy. MYM wrote the first draft of the manuscript and participated in its editing. CJF, RM, and SCF provided feedback on the manuscript. CJF and SCF gained financial support for the project. SCF provided oversight for the entire project.

## Abbreviations

TNBC: Triple negative breast cancer
CA: Carbonic anhydrase
HIF1: Hypoxia Inducible factor 1
CA IX: Carbonic anhydrase isoform IX
CA XII: Carbonic anhydrase isoform XII
CA II: Carbonic anhydrase isoform II
MCT1: Monocarboxylate transporter 1
MCT4: Monocarboxylate transporter 4
vATPase: Vacuolar ATPase
NHE1: Na^+^/H^+^ exchanger
GLUT1: Glucose transporter 1
NBCn1: Na^+^/HCO_3_^−^ co-transporter
PDX: Patient derived tumor graft
USB: Ureido substituted benzene sulfonamide

