## Supplemental Data for "A non-catalytic function of carbonic anhydrase IX contributes to the glycolytic phenotype and pH regulation in human breast cancer cells" (from another version of this preprint: version 2, 2019-02-26)

**Supplemental Figure 1.** Kaplan Meier plots across all breast cancer patients. mRNA from all breast cancer patients (unrestricted analysis) was probed for proton transporters. Panel A) Expression of *ATP6V0A1* gene subunit, a component of the vATPase protein, was probed for using Affimetrix ID 212383_at. Panel B) The gene that encodes the NHE1 protein, *SLC9A1*, was probed for using Affimetrix ID, 209453. Panel C) *SLC16A1* (encodes MCT1 protein) was probed for using Affimetrix ID, 202235_at. Panel D) The *SLC16A3* gene, which encodes the MCT4 protein, was probed for using Affimetrix ID, 202856_at.

**Supplemental Figure 2.** Cell specific protein expression of CAs and transporters in breast cancer cell lines. Cell lines were grown to ~75% confluency and then exposed to normoxic (N), hypoxic conditions (H, 1% $O_2$ for 16 h), or 100 µM desferroxamine mesylate (DFO) for 16 h. After 16 h, the cells were rinsed with PBS, and lysates were extracted using RIPA buffer containing protease inhibitors. Equal concentrations of protein were loaded onto SDS PAGE gels, and then transferred to nitrocellulose for western blot analysis. Membranes were probed for CA expression (CA IX, CA XII and CA II), and transporter expression (V-ATPase, MCT1, MCT4 and GLUT-1). GAPDH was used as a loading control. Comparison between A) MCF10A and UFH-001 cells and B) MCF10A and T47D cells. Data presented represents one of three independent experiments.

**Supplemental Figure 3.** Effect of USB compounds on glucose transport. Panel A) Deoxyglucose uptake was assessed in UFH-001 cells exposed to normoxic conditions or 16 h of hypoxia and in the presence or absence of 5 µM U-$CH_3$ or U-F or U-$NO_2$. Panel B) Deoxyglucose uptake was assessed in T47D cells exposed to normoxic conditions or 16 h of hypoxia and in the presence or absence of 5 µM U-$CH_3$ or U-F or U-$NO_2$. n = 3 for all data sets.

Supporting Figure 1

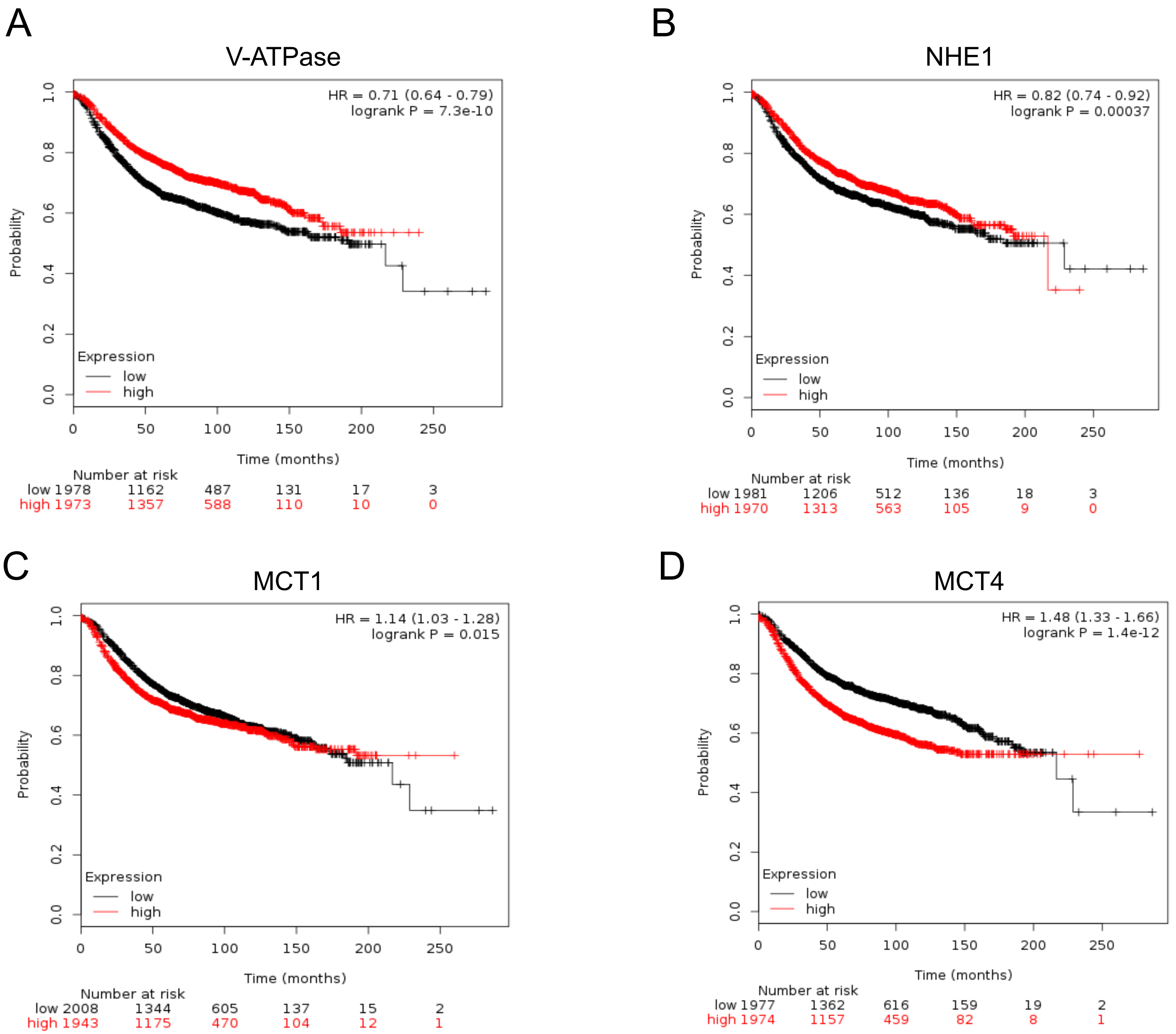

Supporting Figure 2

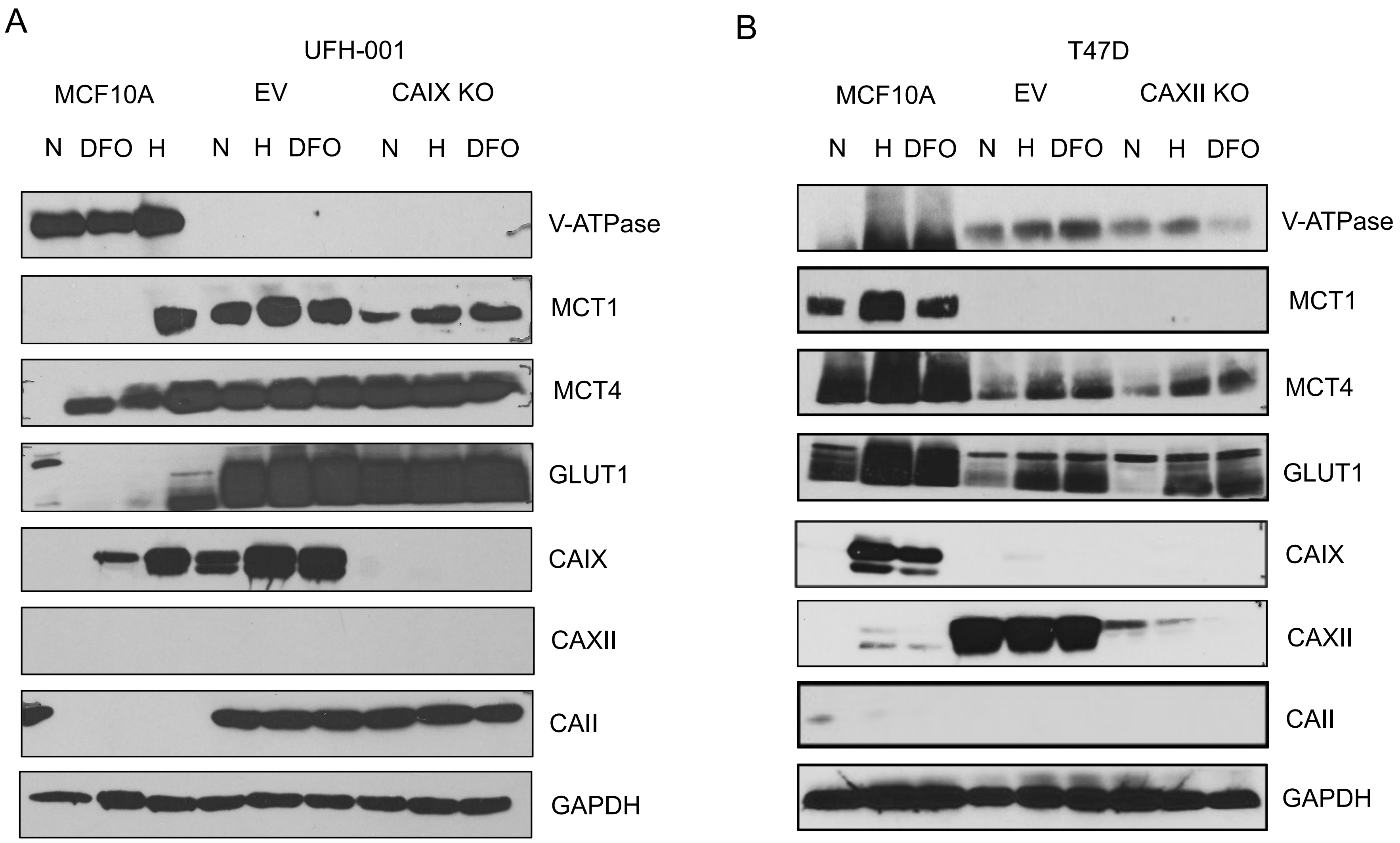

### Supporting Figure 3

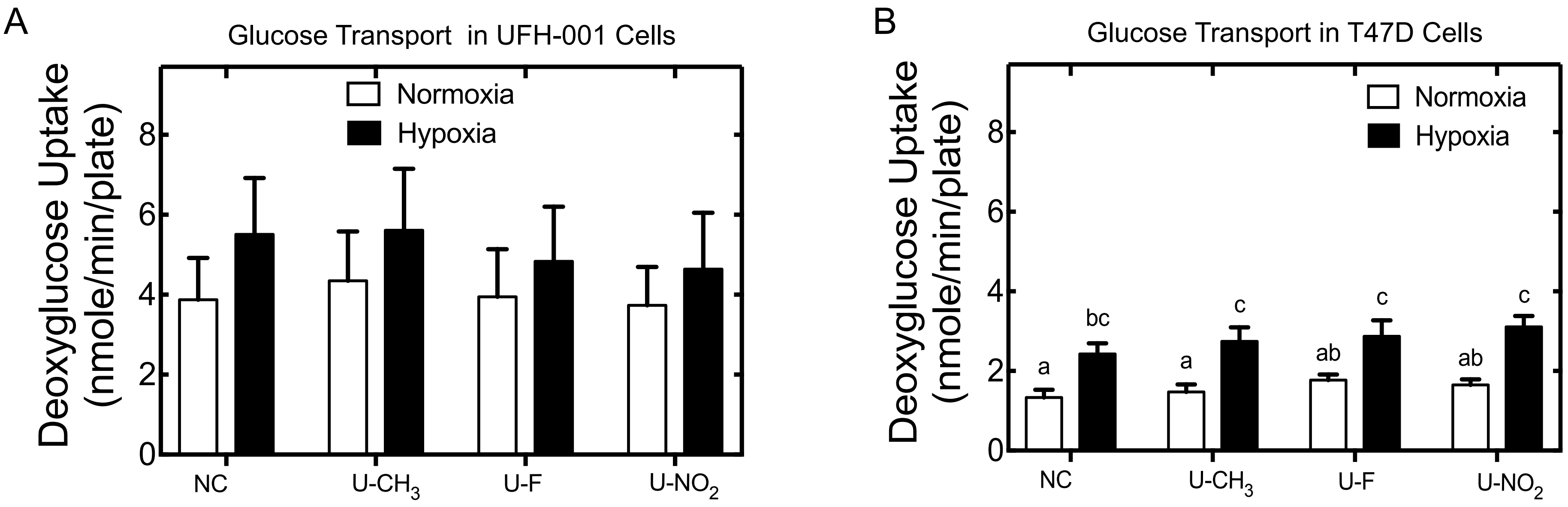
